# Comparative analysis of sequential and thermodynamic features of pre-miRNA in insects with various organisms and applying XGBoost for one-vs-rest binary classification

**DOI:** 10.1101/2025.09.17.676766

**Authors:** Adhiraj Nath, Utpal Bora

## Abstract

MicroRNAs are found to regulate various biological processes which are produced from precursor microRNA. As the length of such microRNA are small, homology-based searching is not very useful. Hence, various machine learning based tools have been designed for prediction of such hairpin loops using various thermodynamic and sequential features. In this research, we discuss about the comparative statistical analysis of various features used the in development of machine learning based predictive tools. The sequence features of insect precursor microRNA were compared with precursor microRNA of other available organisms. We initially established that features such as Length, GC content, Minimum Free Energy (MFE) of folding, etc., differs in insects as compared to other organisms using Kolmogorov-Smirnov (KS) test. We further trained a predictive model for one-vs-rest binary classification using XGBoost between insects, human, monocots, aves, ruminants, sauria, dogs and rodents. We performed PCA and retained 14 principal components for classification using cumulative explained variance. Various parameters of XGBoost was tuned with 5-fold CV and the parameter values with highest CV score were considered. We used independent held-out data test the models. The accuracy of insect, monocots, rodents, human, ruminants, sauria, aves and dogs was found to be 0.8549, 0.8626, 0.6835, 0.7005, 0.8875, 0.6972, 0.7591 and 0.6588 respectively. This shows that ancestral lineage specific ML models can be developed for detection of precursor microRNA for different classes of organism.

## 1. Background

Precursor microRNA (pre-miRNA) are the non-coding RNA hairpin loops which is cleaved by Drosha to produce microRNA (miRNA) ^1,2^. Multiple miRNAs can be produced from a single pre-miRNA for which characterization and identification of pre-miRNA has been of great importance. miRNA has been found to regulate gene expression of various biological processes such as development, cell proliferation, cell differentiation, apoptosis, transposon silencing, and antiviral defense ^3–6^. In insects, changes in miRNA expression profile have been observed in various biological processes such as metamorphosis, reproduction, immune response, etc. ^7–13^. miRNAs are believed to be conserved although they target diverse genes. They are believed to be similar across all the species. ^14–16^

Various tools are designed to predict pre-miRNAs as they give rise to mature miRNA. These data are downloaded from miRBase which contains collection of pre-miRNAs and their corresponding miRNAs of various organisms ^17^. It currently holds miRNAs from 271 organisms. Features such as nucleotide bases, length of the sequence, GC content of pre-miRNAs are used to train machine learning classifiers to predict a true pre-miRNA ^18–30^. Deep learning methods were also carried out for detecting pre-miRNA hairpin loop in COVID. ^31^

However, most existing tools are either general-purpose or tailored to a single organism or taxonomic group, and they often assume that pre-miRNA features are conserved across species. This assumption may not hold for phylogenetically distant groups such as insects, which are known to have unique regulatory networks and ecological specializations. There is a growing need to develop organism-aware or lineage-specific models to improve the accuracy and biological relevance of miRNA prediction. ^32,33^.

In this work, we analyzed the pre-miRNA sequences of insect pre-miRNA from miRbase and performed comparative statistical analysis with other available organisms. We initially established that features such as Length, GC content, MFE, etc., differs in insects as compared to other organisms. We further trained a predictive model for classification using XGBoost between insects, human, monocots, aves, ruminants, sauria, dogs and rodents.

## 2. Methods

### 2.1. Data Collection and pre-processing

We collected pre-miRNA sequences of insects, human, monocots, aves, ruminants, sauria, dogs and rodents from miRBase ^17^ and labelled them for comparison. The secondary structure was calculated using RNAfold software from ViennaRNA package. The fasta header, nucleotide sequence, MFE score and secondary structure for each pre-miRNA sequence was converted into tabular format using in-house python script.

### 2.2. Hypothesis testing

The null hypothesis was:

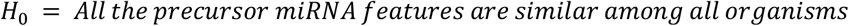

Our alternate hypothesis states that insect pre-miRNAs are different in many aspects which are routinely used in ML (machine learning) tools, i.e.

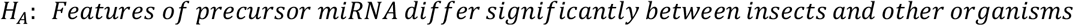

To determine if the normality of distribution Shapiro-Wilk test ^34^ was performed given in equation 1.

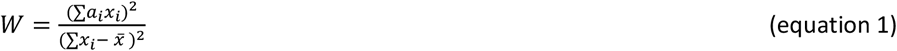

Where: **x**_***i***_: are the ordered sample values.

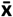: is the sample mean.

**a**_***i***_: are coefficients that depend on the sample size n

The results suggested that the data was not normally distributed and hence, we performed two-sample Kolmogorov-Smirnov (KS) tests given in equation 2, to compare the distributions of 57 pre-miRNA features between insects and each of seven other organisms (aves, human, mammalia, monocots, rodent, rumin, sauria), resulting in 399 comparisons.

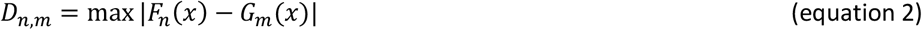

Where: F_n_(x) is the theoretical CDF and G_m_(x) is the empirical CDF.

The significance level for all statistical tests was set at α = 0.05. Additionally, to account for multiple comparisons, a Bonferroni correction was applied to adjust the p-values and minimize the risk of type I errors.

In order to apply ml algorithms, we also performed Mann-Whitney U tests to assess median differences, and Levene’s tests to evaluate variance equality.

### 2.3. Feature engineering

We used dimensionality reduction techniques to identify the most informative features for training. Initially, we calculated Pearson’s correlation coefficients between all feature pairs to detect multicollinearity. For each pair of highly correlated features, one representative feature was retained while the other was removed to eliminate redundancy and reduce overfitting risk. Following this filtering step, we applied Principal Component Analysis (PCA) and computed the sum of squared loadings for each feature to assess its contribution to the principal components. We then selected the minimal set of features that collectively accounted for 95% of the total variance in the dataset also known as Cumulative Explained Variance ^35,36^, ensuring both compactness and informativeness of the feature set.

### 2.4. Training with XGBoost

A one-vs-rest binary classification approach was adopted to distinguish each organism from the rest. For each target organism, all available sequences were treated as positive samples. An equal number of negative samples were randomly drawn without replacement from the pool of remaining organisms to maintain class balance. This procedure ensured that each binary classifier was trained on a balanced dataset of equal positive and negative examples. We kept 80% of the data for training (X_train) and used 20% of the data for testing (X_test). We used XGBoost classifier to train the data for classification. The algorithm of XGBoost constructs multiple CART models in parallel which effectively improves the computation speed. Second-order Taylor formula is used to optimize by model by calculating the error value between the predicted and true value ^37^. It can further handle missing feature values by internally imputing it and hence does not require feature standardization ^38^. It has hence been used in the estimation and classification biological data ^39,40^. It is based on minimising the loss function and regularization, *L*^*(t)*^ which mathematically it can be written as:

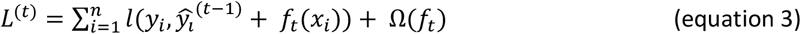

Where *l* measures the difference between prediction 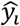 and target *y*_*i*_ in the *i*th instance at iteration *t. f*_*t*_ is an independent tree for given input *x*_*i*_. Ω(*f*_*t*_) works has a penalty function.

We used 5 fold CV while optimising the following parameters of the XGBoost package in Scikit-learn:

- **lambda**: L2 regularization range from 1e-3 to 10
- **alpha**: L1 regularization range from 1e-3 to 10.
- **colsample_bytree**: Subsample ratio of columns during construction of each tree, ranges from 0.3 to 1.0
- **subsample**: Ratio of training instances, ranges from 0.4 to 1
- **learning_rate**: Step size at each iteration while moving towards minimum of loss function, ranges from 0.001 to 0.2
- **n_estimators**: Number of trees, ranges from 50 to 400
- **max_depth**: Max depth of a tree, ranges from 5 to 17
- **min_child_weight**: Minimum instances needed to be in each node, ranges from 1 to 300

### 2.5. Performance Analysis

To assess the generalizability of each classifier, the hyperparameter tuning and cross-validation were carried out exclusively on the training data and the final model was evaluated on the held-out test set.

The best parameters were chosen and the models were evaluated on X_test dataset to check their efficiency. During performance evaluation, we considered True dataset to be the group that is being evaluated and False dataset to be the collection of all other groups in each case. The performance was calculated based on the following classical classification measures: sensitivity 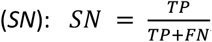, specificity 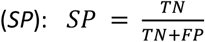, Accuracy 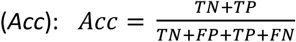, precision 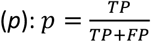, harmonic mean of sensitivity and precision 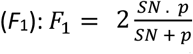 and Matthew’s correlation coefficient (*MCC*): 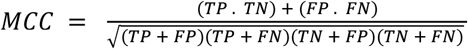, where *TP, TN, FP* and *FN* are the number of true-positive, false-positive and false-negative classifications, respectively. For given false positive rate (α) and true positive rate (1 − β) at different threshold values, the AUC-ROC was computed as: 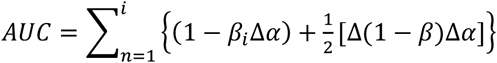 where Δ(1−β)=(1−β_i_)−(1−β_i−1_) and Δα=α_i_−α_i−1_ and i = 1, 2, …, m (number of test data points) ^33^. The workflow of the methods is given in Figure 1.

**Figure 1.**
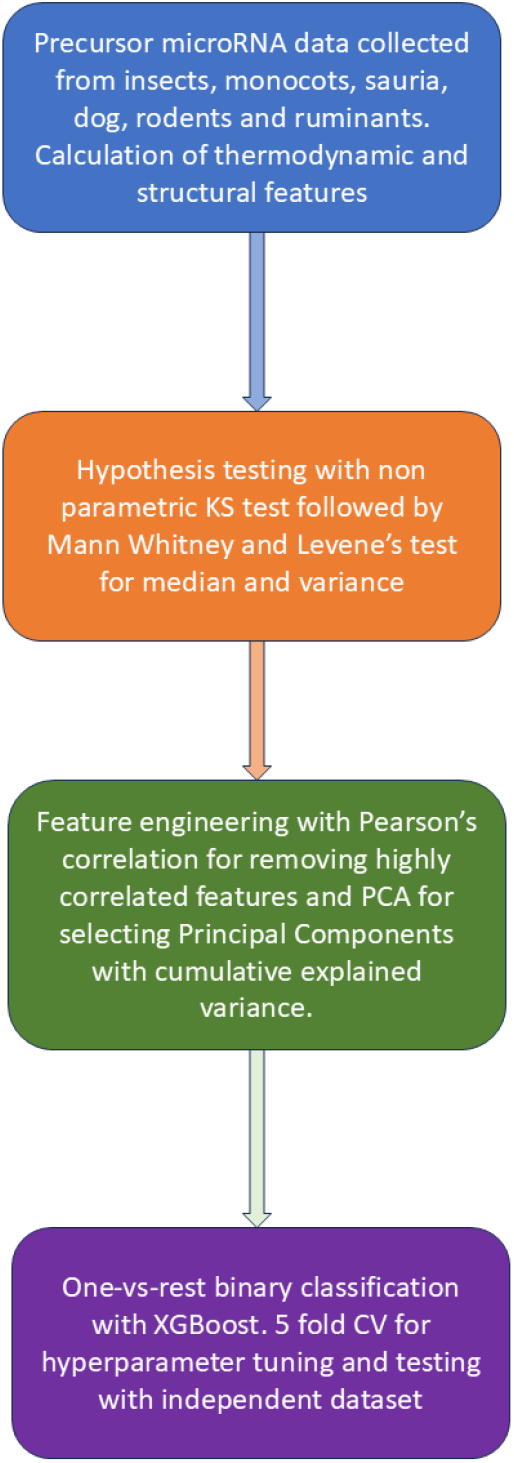
Workflow for XGBoost training.

## 3. Results

### 3.1. Data pre-processing

A total of 5541 sequences were collected for the analysis as given in Table 1.

**Table 1:**
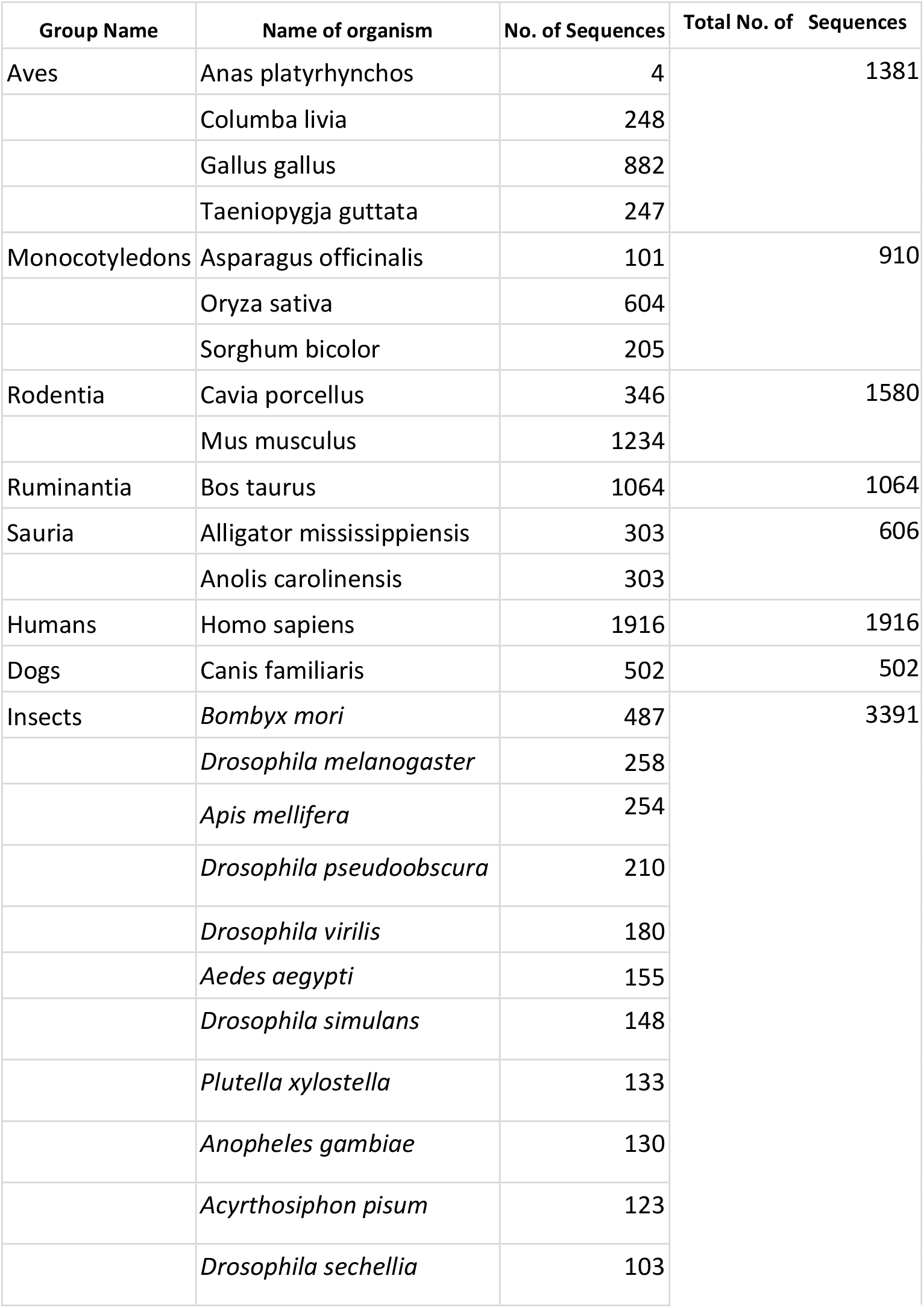
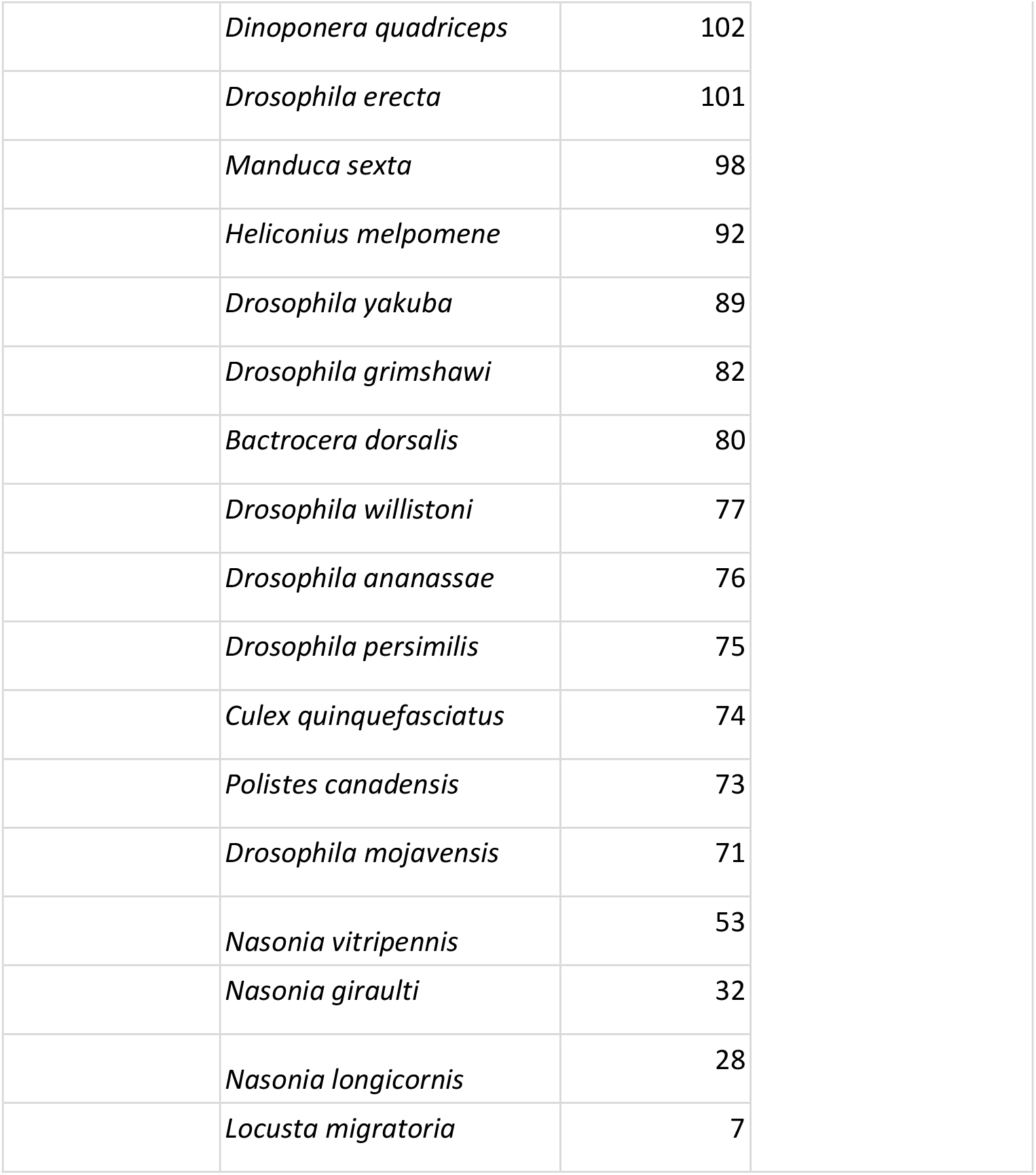
Total sequences collected for the analysis.

| Group Name | Name of organism | No. of Sequences | Total No. of Sequences |
| --- | --- | --- | --- |
| Aves | Anas platyrhynchos | 4 | 1381 |
|  | Columba livia | 248 |  |
|  | Gallus gallus | 882 |  |
|  | Taeniopygia guttata | 247 |  |
| Monocotyledons | Asparagus officinalis | 101 | 910 |
|  | Oryza sativa | 604 |  |
|  | Sorghum bicolor | 205 |  |
| Rodentia | Cavia porcellus | 346 | 1580 |
|  | Mus musculus | 1234 |  |
| Ruminantia | Bos taurus | 1064 | 1064 |
| Sauria | Alligator mississippiensis | 303 | 606 |
|  | Anolis carolinensis | 303 |  |
| Humans | Homo sapiens | 1916 | 1916 |
| Dogs | Canis familiaris | 502 | 502 |
| Insects | *Bombyx mori* | 487 | 3391 |
|  | *Drosophila melanogaster* | 258 |  |
|  | *Apis mellifera* | 254 |  |
|  | *Drosophila pseudoobscura* | 210 |  |
|  | *Drosophila virilis* | 180 |  |
|  | *Aedes aegypti* | 155 |  |
|  | *Drosophila simulans* | 148 |  |
|  | *Plutella xylostella* | 133 |  |
|  | *Anopheles gambiae* | 130 |  |
|  | *Acyrtosiphon pisum* | 123 |  |
|  | *Drosophila sechellia* | 103 |  |

Table 1: Total sequences collected for the analysis.
|  |  |  |
| --- | --- | --- |
|  | *Dinoponera quadriceps* | 102 |
|  | *Drosophila erecta* | 101 |
|  | *Manduca sexta* | 98 |
|  | *Heliconius melpomene* | 92 |
|  | *Drosophila yakuba* | 89 |
|  | *Drosophila grimshawi* | 82 |
|  | *Bactrocera dorsalis* | 80 |
|  | *Drosophila willistoni* | 77 |
|  | *Drosophila ananassae* | 76 |
|  | *Drosophila persimilis* | 75 |
|  | *Culex quinquefasciatus* | 74 |
|  | *Polistes canadensis* | 73 |
|  | *Drosophila mojavenensis* | 71 |
|  | *Nasonia vitripennis* | 53 |
|  | *Nasonia giraulti* | 32 |
|  | *Nasonia longicornis* | 28 |
|  | *Locusta migratoria* | 7 |

### 3.2. Parameter Calculation

The parameters used for the classification is given in Table 2. A total of 57 parameters were calculated out of which 16 were dinucleotide counts and dinucleotide percentage counts, 4 nucleotide counts and their percentage counts, 2 base pair counts, base propensity, Shannon entropy, etc.

**Table 2.:**
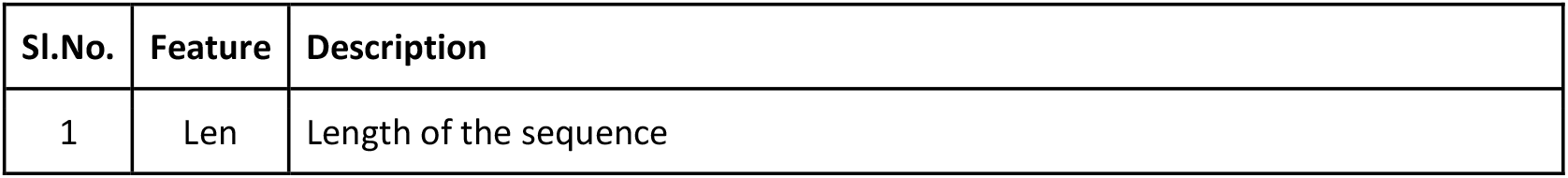
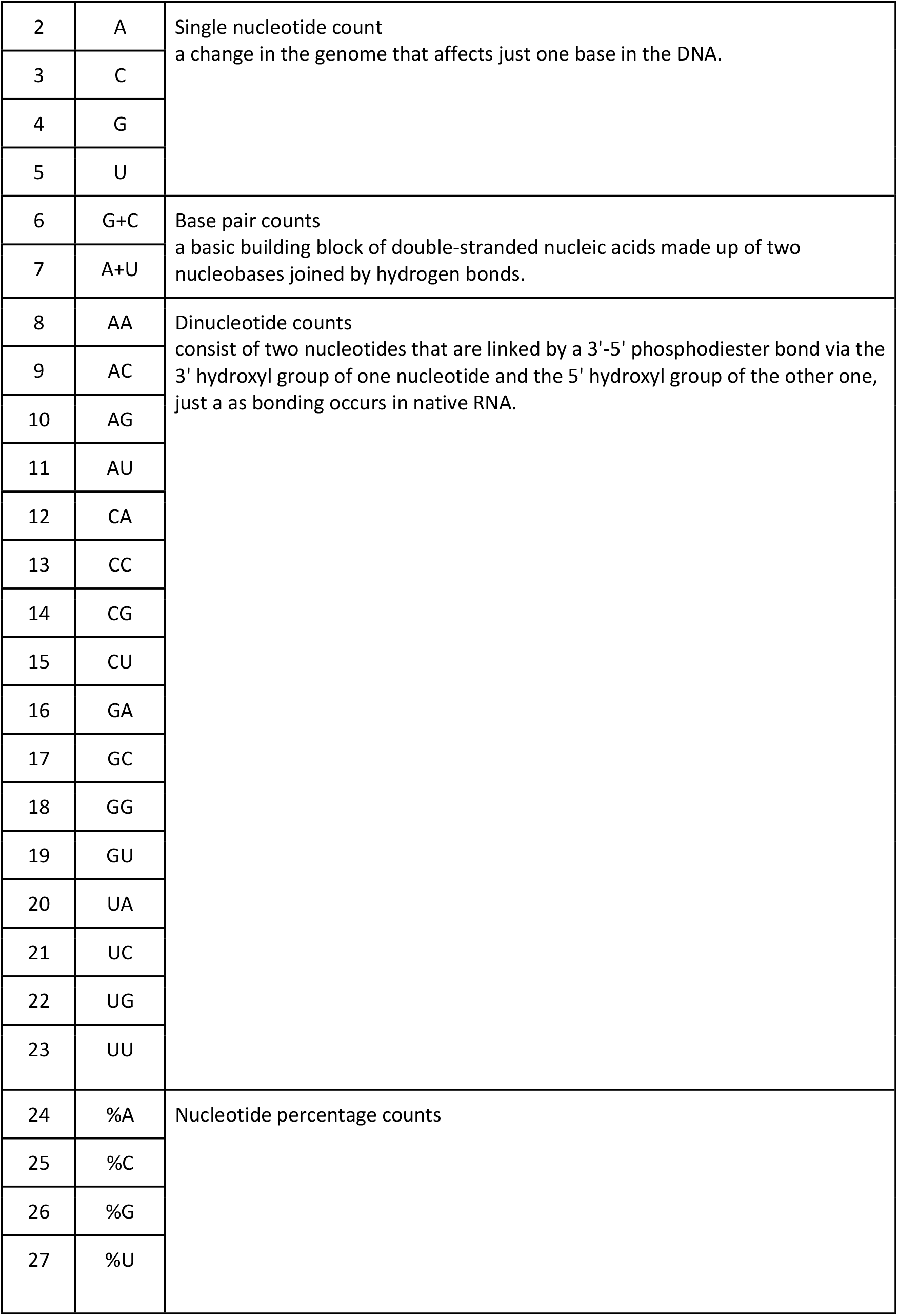
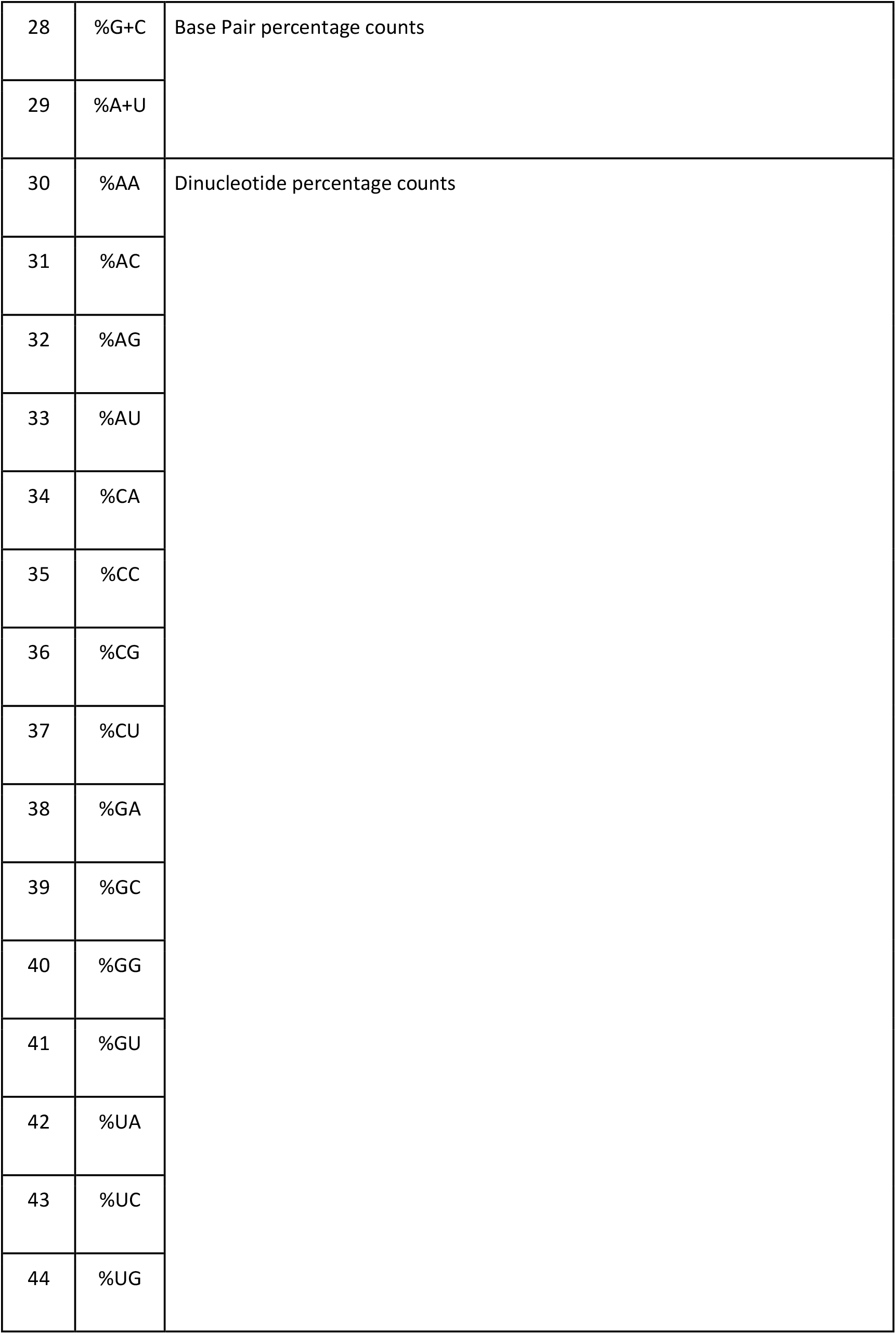
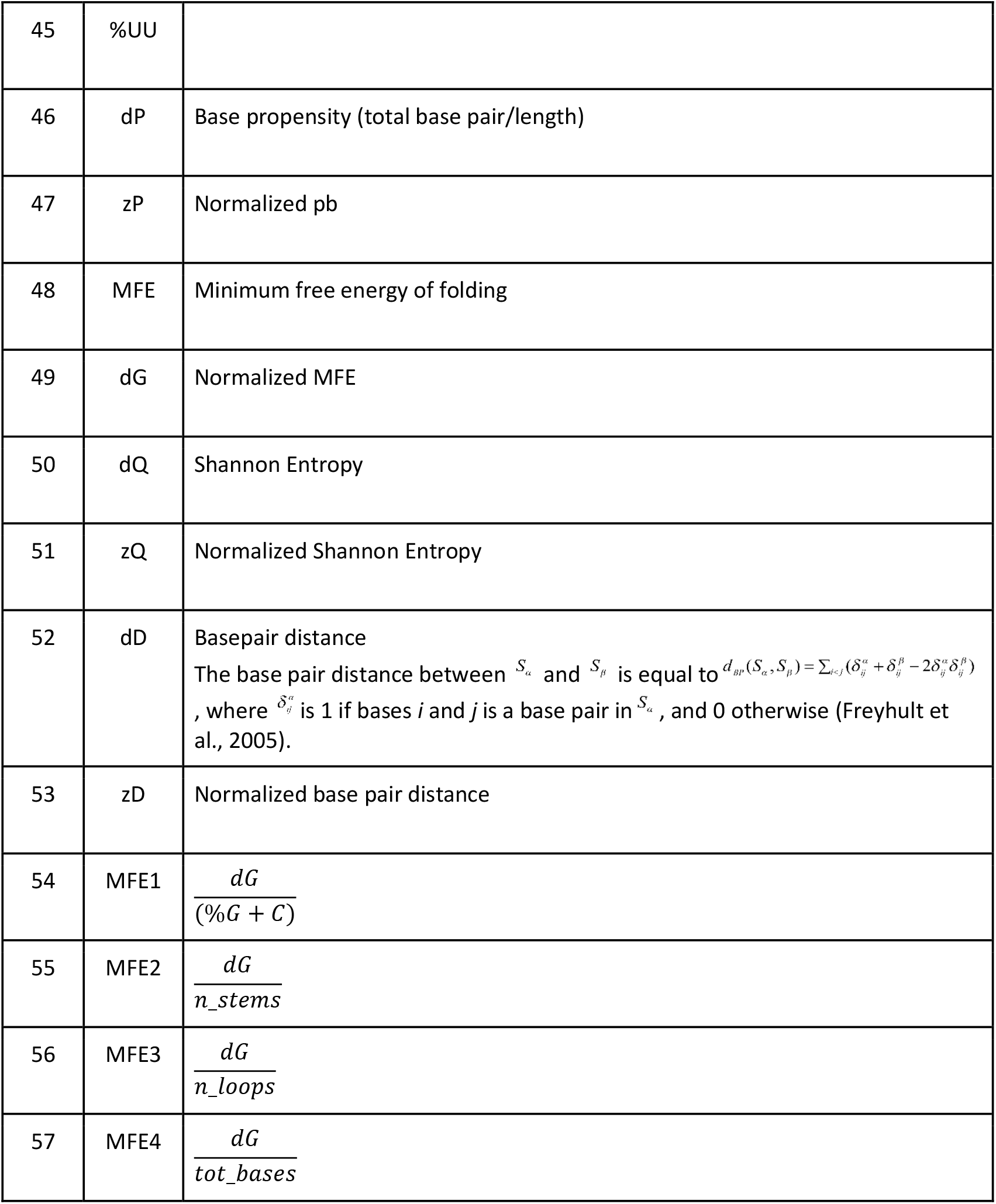
Parameters calculated for feature extraction.

| Sl.No. | Feature | Description |
| --- | --- | --- |
| 1 | Len | Length of the sequence |
| 2 | A | Single nucleotide count<br>a change in the genome that affects just one base in the DNA. |
| 3 | C |  |
| 4 | G |  |
| 5 | U |  |
| 6 | G+C | Base pair counts<br>a basic building block of double-stranded nucleic acids made up of two nucleobases joined by hydrogen bonds. |
| 7 | A+U |  |
| 8 | AA | Dinucleotide counts<br>consist of two nucleotides that are linked by a 3'-5' phosphodiester bond via the 3' hydroxyl group of one nucleotide and the 5' hydroxyl group of the other one, just as bonding occurs in native RNA. |
| 9 | AC |  |
| 10 | AG |  |
| 11 | AU |  |
| 12 | CA |  |
| 13 | CC |  |
| 14 | CG |  |
| 15 | CU |  |
| 16 | GA |  |
| 17 | GC |  |
| 18 | GG |  |
| 19 | GU |  |
| 20 | UA |  |
| 21 | UC |  |
| 22 | UG |  |
| 23 | UU |  |
| 24 | %A | Nucleotide percentage counts |
| 25 | %C |  |
| 26 | %G |  |
| 27 | %U |  |
| 28 | %G+C | Base Pair percentage counts |
| 29 | %A+U |  |
| 30 | %AA | Dinucleotide percentage counts |
| 31 | %AC |  |
| 32 | %AG |  |
| 33 | %AU |  |
| 34 | %CA |  |
| 35 | %CC |  |
| 36 | %CG |  |
| 37 | %CU |  |
| 38 | %GA |  |
| 39 | %GC |  |
| 40 | %GG |  |
| 41 | %GU |  |
| 42 | %UA |  |
| 43 | %UC |  |
| 44 | %UG |  |
| 45 | %UU |  |
| 46 | dP | Base propensity (total base pair/length) |
| 47 | zP | Normalized pb |
| 48 | MFE | Minimum free energy of folding |
| 49 | dG | Normalized MFE |
| 50 | dQ | Shannon Entropy |
| 51 | zQ | Normalized Shannon Entropy |
| 52 | dD | Basepair distance<br>The base pair distance between $S_\alpha$ and $S_\beta$ is equal to $d_{BP}(S_\alpha, S_\beta) = \sum_{i < j} (\delta_{ij}^\alpha + \delta_{ij}^\beta - 2\delta_{ij}^\alpha \delta_{ij}^\beta)$, where $\delta_{ij}^\alpha$ is 1 if bases $i$ and $j$ is a base pair in $S_\alpha$, and 0 otherwise (Freyhult et al., 2005). |
| 53 | zD | Normalized base pair distance |
| 54 | MFE1 | $\frac{dG}{(\%G + C)}$ |
| 55 | MFE2 | $\frac{dG}{n\_stems}$ |
| 56 | MFE3 | $\frac{dG}{n\_loops}$ |
| 57 | MFE4 | $\frac{dG}{tot\_bases}$ |

### 3.3. Hypothesis Testing

The Shapiro-Wilk test results indicate that the parameters are not normally distributed, suggesting the use of the Kolmogorov–Smirnov (KS) test. A Bonferroni-corrected alpha threshold of 0.000125 was applied to the KS test, which identified 335 combinations as significantly different across various organism classes. In the remaining 64 combinations, no single parameter was common to all organisms, suggesting that the null hypothesis does not hold. Furthermore, the Mann–Whitney U test and Levene’s test showed that all 57 parameters exhibited significant differences in median and variance between at least one pair of organisms. The test results are provided in Supplementary Material 1. Figure 2 illustrates the comparison of selected parameters—including length, %G+C content, and minimum free energy (dG)—across 500 randomly sampled pre-miRNA sequences from each organism for visual clarity.

**Figure 2:**
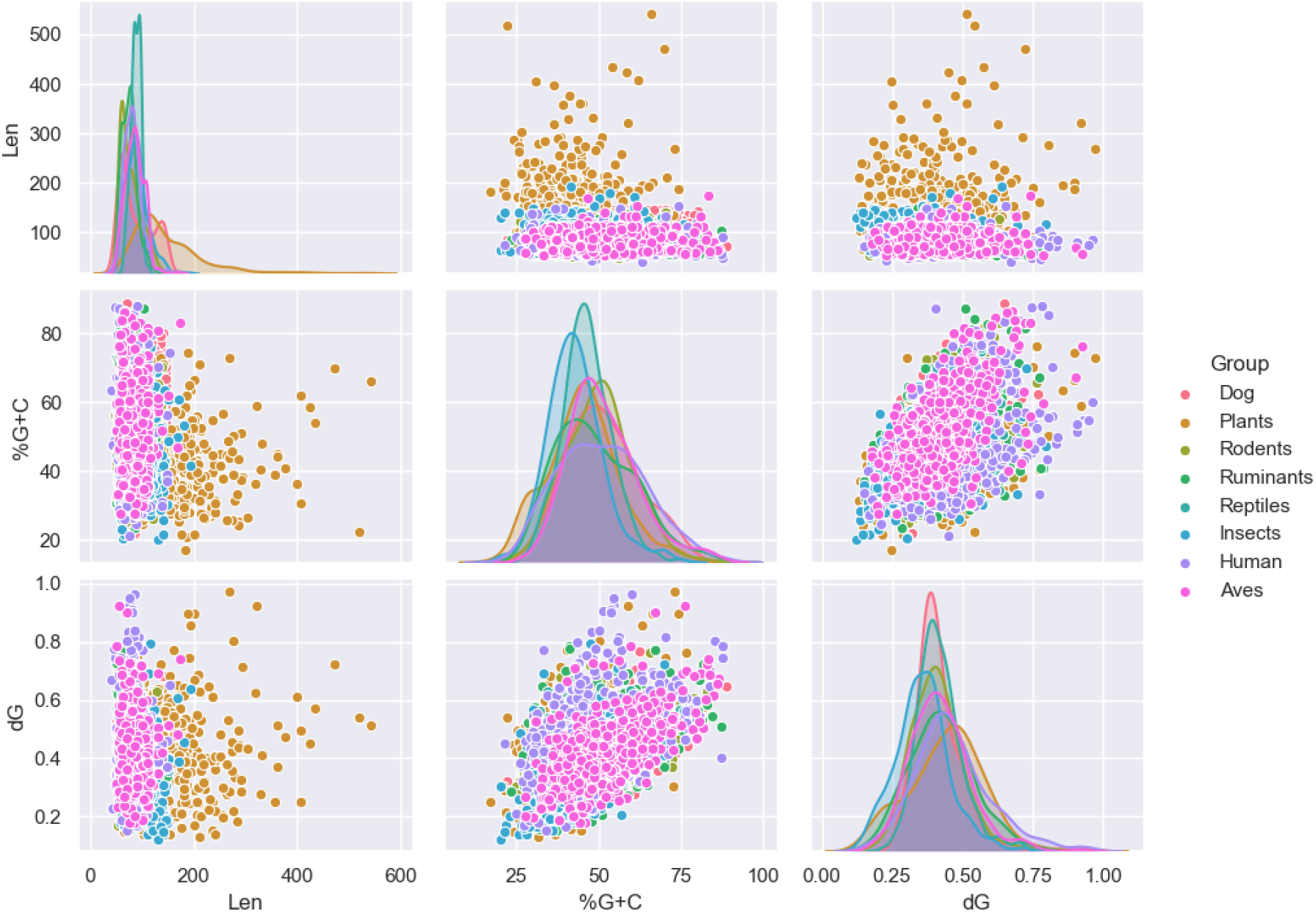
Comparison of insect pre-miRNA with other class of organisms. The comparation of various features of 500 randomly sampled pre-miRNA from insects, human, monocots, aves, ruminants, sauria, dogs and rodents are is shown in the pair-plot scatter diagrams. Features such as GC percentage (%G+C), Length (Len) and dG (MFE/Length) were considered out of 57 features mentioned below. Multivariate gaussian distribution plot is given in the diagonals.

### 3.4. Feature Engineering

To reduce redundancy, only one feature from each group of highly correlated variables was retained. As a result, the following features were removed due to high correlation: ‘AU’, ‘ND’, ‘A+U’, ‘G’, ‘G+C’, ‘A’, ‘%GC’, ‘%GG’, ‘%CC’, ‘%UU’, ‘mfe’, ‘D’, ‘AA’, ‘CC’, ‘U’, ‘%UA’, ‘NQ’, ‘pb’, ‘MFE3’, ‘%CG’, ‘%A+U’, ‘UU’ and ‘%G+C’. The results of the PCA are presented in Table 3. Based on the cumulative explained variance, 14 principal components were selected, capturing 95% of the total variance.

**Table 3:**
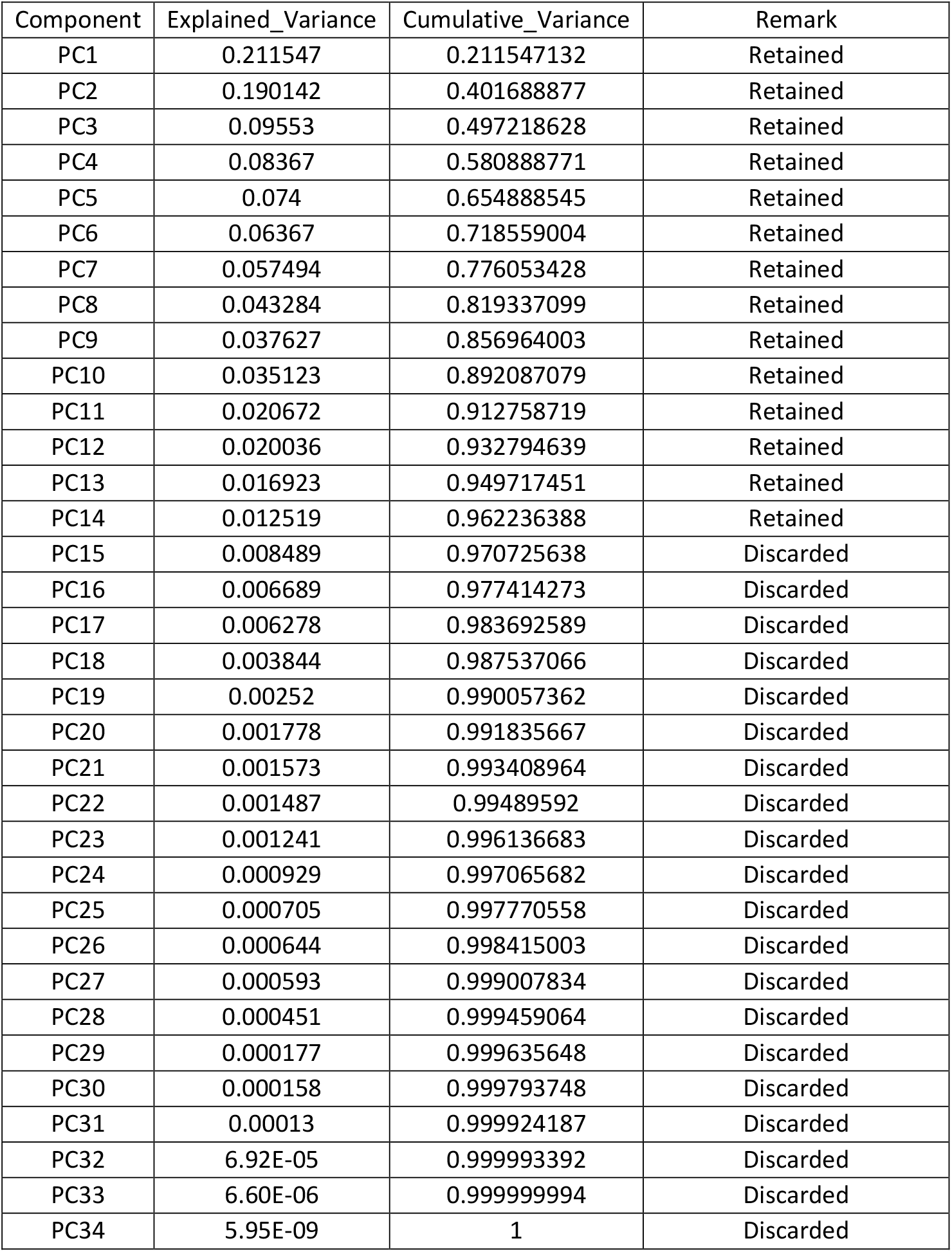
Principal Component Analysis (PCA) results showing the explained variance and cumulative variance for each component. Based on the criterion of capturing 95% of the cumulative variance, the first 14 principal components were retained for further analysis, while the remaining components were discarded.

The cumulative explained variance by each principal component is also illustrated in Figure 3. Based on this plot, the first 14 components were selected, as they collectively account for approximately 95% of the total variance.

**Figure 3:**
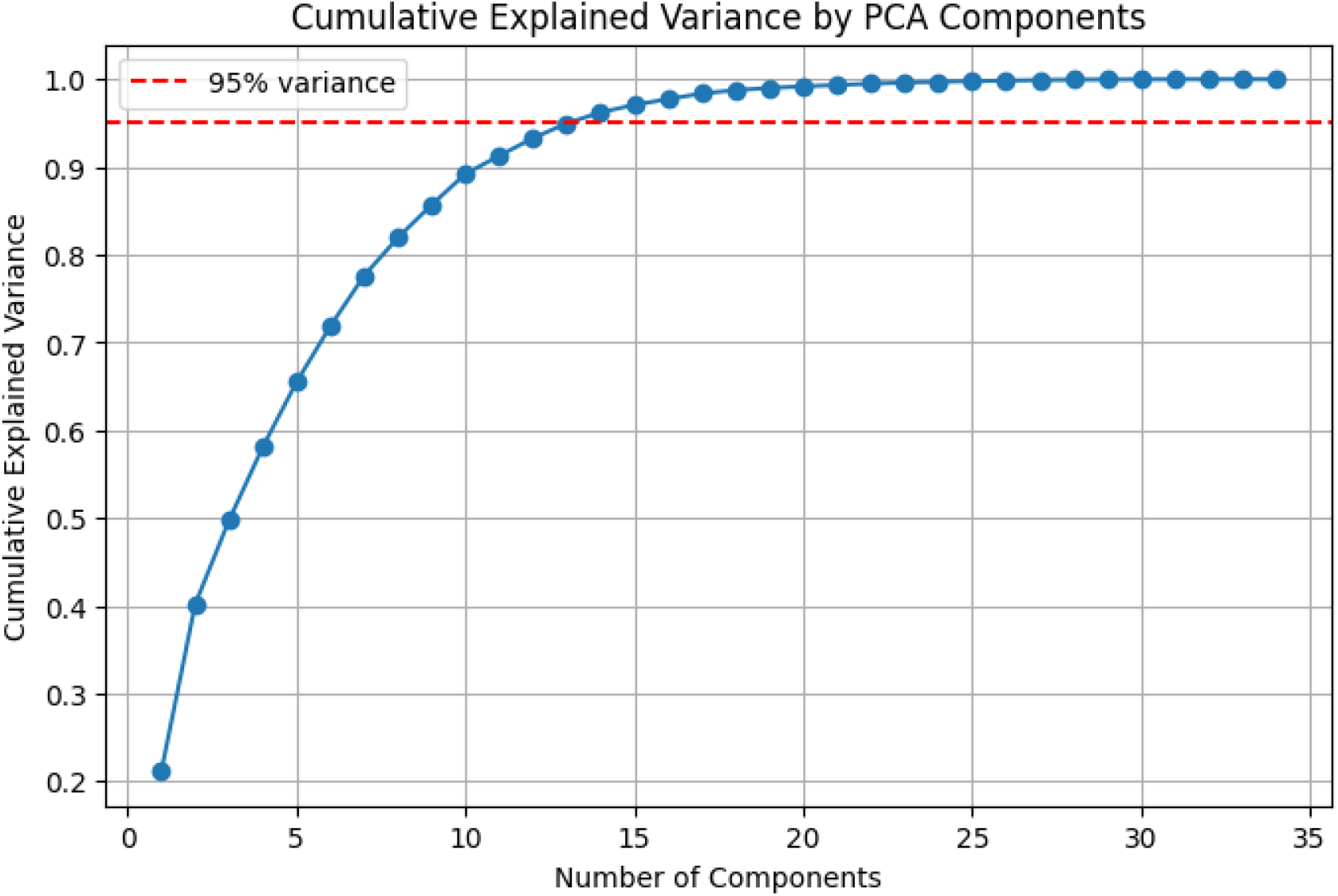
Cumulative explained variance of principal components derived from PCA. The first 14 components, accounting for 95% of the total variance, were retained for downstream analysis. The elbow in the curve indicates the point of diminishing returns for additional components.

### 3.5. Training with XGBoost

The classification model for each organism was trained and evaluated using a random search cross-validation strategy on 80% of the training dataset followed by testing on 20% of the independent hold-out dataset. Table 4 summarizes the best cross-validation (CV) accuracy along with test accuracy, precision, recall, and F1-score. Detailed results of the hyperparameter tuning process—including fit times, fold-wise test scores, selected hyperparameter values, and model ranking—for each organism are provided in Supplementary Material 2.

**Table 4:** Performance metrics of the classification model for each organism. The table includes the best cross-validation (CV) accuracy and corresponding test set metrics: accuracy, precision, recall, and F1-score. Higher scores for monocots, insects, and sauria suggest more distinct or learnable features, while lower scores in other groups indicate potential classification challenges.

| Organism | Best CV Accuracy | Test Accuracy | Test Precision | Test Recall | Test F1 |
| --- | --- | --- | --- | --- | --- |
| aves | 0.68704097 | 0.658844765 | 0.650684932 | 0.685920578 | 0.667838 |
| human | 0.706591391 | 0.700520833 | 0.702631579 | 0.6953125 | 0.698953 |
| dogs | 0.693330745 | 0.668316832 | 0.697674419 | 0.594059406 | 0.641711 |
| monocots | 0.836522619 | 0.862637363 | 0.888235294 | 0.82967033 | 0.857955 |
| rodent | 0.6890823 | 0.683544304 | 0.669590643 | 0.724683544 | 0.696049 |
| rumin | 0.651607728 | 0.704225352 | 0.700460829 | 0.713615023 | 0.706977 |
| sauria | 0.820303403 | 0.795081967 | 0.78125 | 0.819672131 | 0.8 |
| insect | 0.832038193 | 0.854933726 | 0.866261398 | 0.839469809 | 0.852655 |

The best hyperparameter values identified during model tuning for each organism are summarized in Table 5. These include values for tree-specific parameters (e.g., max_depth, min_child_weight), learning rate, regularization parameters (reg_alpha, reg_lambda), and sampling parameters (colsample_bytree, subsample).

**Table 5:**
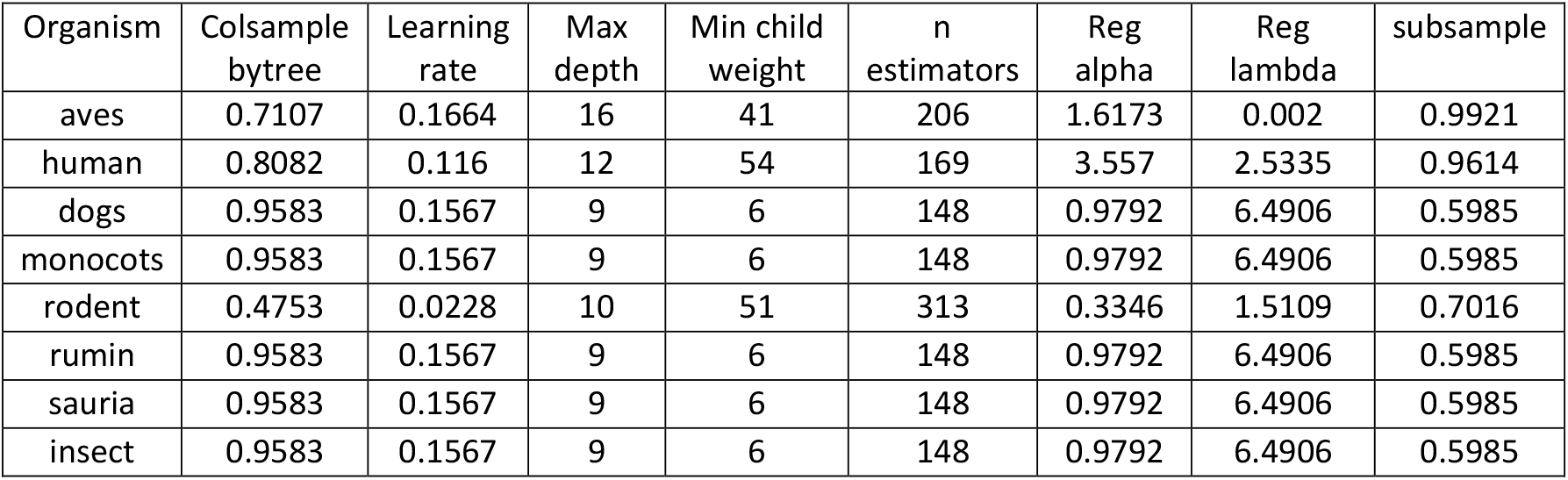
Best-performing hyperparameter values for each organism as identified through cross-validation. Parameters include colsample_bytree, learning_rate, max_depth, min_child_weight, n_estimators, reg_alpha, reg_lambda, and subsample.

### 3.6. Performance Evaluation

Various performance measures for each group of organisms is given in Table 6. The accuracy of insect, monocots, rodents, human, ruminants, sauria, aves and dogs was found to be 0.8549, 0.8626, 0.6835, 0.7005, 0.8875, 0.6972, 0.7591 and 0.6588 respectively. Specificity was found to be 0.8704 for insects, 0.8956 for monocots, 0.6424 for rodents, 0.7057 for human, 0.7042 for ruminants, 0.7705 for sauria, 0.6318 for aves and 0.7426 for dogs. The F1 score of insect, monocots, rodents, human, ruminants, sauria, aves and dogs was found to be 0.8572, 0.858, 0.696, 0.699, 0.695, 0.8, 0.6678 and 0.6415 respectively. Sensitivity was found to be 0.8395 for insects, 0.8297 for monocots, 0.7247 for rodents, 0.6953 for human, 0.6901 for ruminants, 0.8197 for sauria, 0.6859 for aves and 0.5941 for dogs. The MCC score of insect, monocots, rodents, human, ruminants, sauria, aves and dogs was found to be 0.7102, 0.7269, 0.3683, 0.4011, 0.3944, 0.5909, 0.3182 and 0.3404 respectively. This has also been graphically shown in Figure 4. The radar plot provides a comparative overview of model performance across organisms, highlighting strengths and weaknesses in different metrics. Notably, organisms such as Monocots and Insects exhibit consistently higher scores, while Aves and Rodents show comparatively lower performance across several metrics.

**Table 6.**
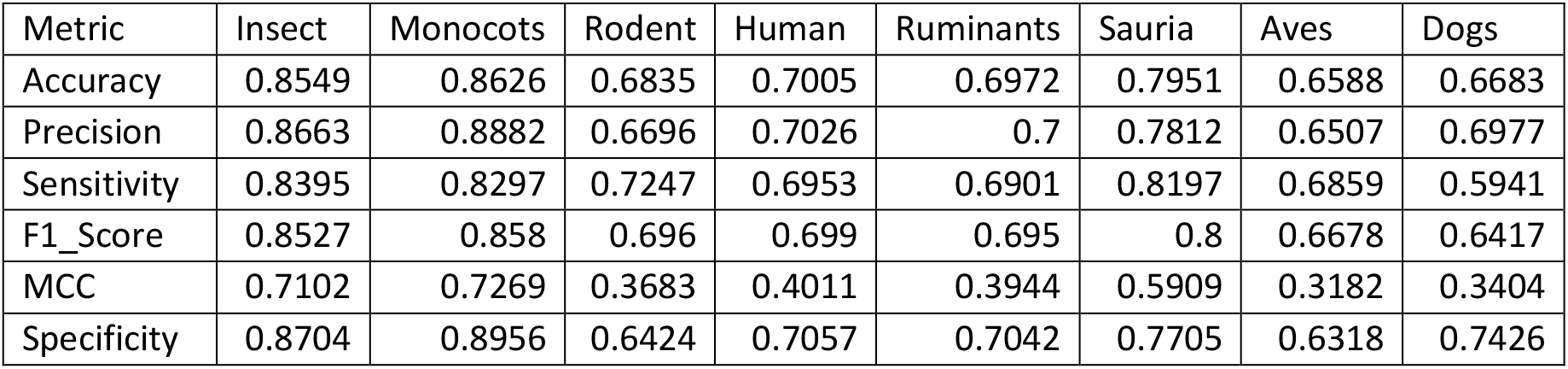
Performance metrics of classification models evaluated across eight different organism groups. Metrics include Accuracy, Precision, Sensitivity, F1 Score, Matthews Correlation Coefficient (MCC), and Specificity. These values reflect the models’ ability to generalize and perform consistently across diverse biological taxa.

| Metric | Insect | Monocots | Rodent | Human | Ruminants | Sauria | Aves | Dogs |
| --- | --- | --- | --- | --- | --- | --- | --- | --- |
| Accuracy | 0.8549 | 0.8626 | 0.6835 | 0.7005 | 0.6972 | 0.7951 | 0.6588 | 0.6683 |
| Precision | 0.8663 | 0.8882 | 0.6696 | 0.7026 | 0.7 | 0.7812 | 0.6507 | 0.6977 |
| Sensitivity | 0.8395 | 0.8297 | 0.7247 | 0.6953 | 0.6901 | 0.8197 | 0.6859 | 0.5941 |
| F1_Score | 0.8527 | 0.858 | 0.696 | 0.699 | 0.695 | 0.8 | 0.6678 | 0.6417 |
| MCC | 0.7102 | 0.7269 | 0.3683 | 0.4011 | 0.3944 | 0.5909 | 0.3182 | 0.3404 |
| Specificity | 0.8704 | 0.8956 | 0.6424 | 0.7057 | 0.7042 | 0.7705 | 0.6318 | 0.7426 |

**Figure 4:**
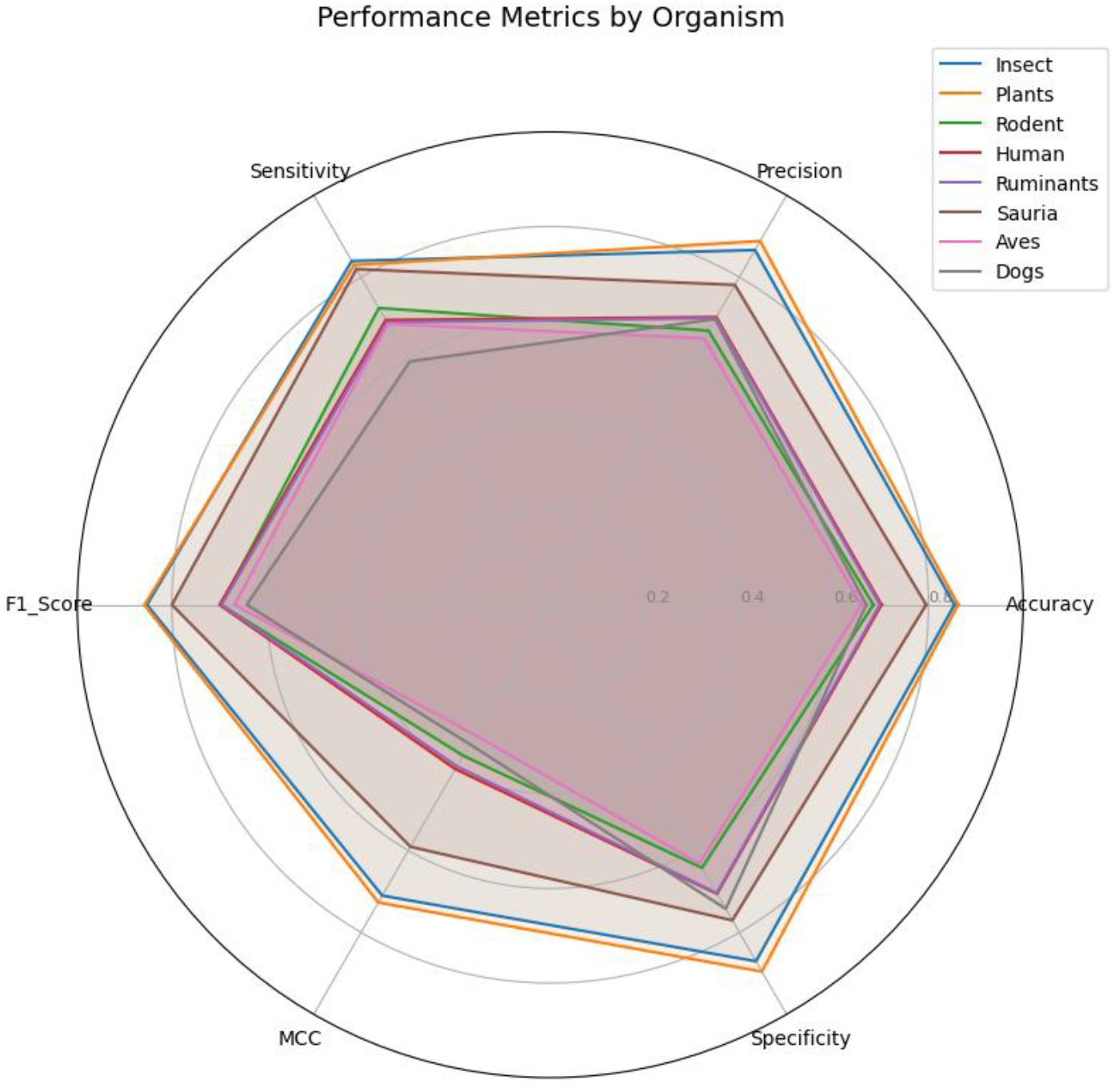
Radar plot illustrating the performance of classification models across different organisms using six evaluation metrics: Accuracy, Precision, Sensitivity, F1 Score, Matthews Correlation Coefficient (MCC), and Specificity. Each line represents a different organism, allowing for comparative visualization of model strengths and weaknesses across taxa.

The ROC-AUC is given in Figure 5. The AUC was found to be 0.92 for Insects, 0.93 for Monocots, 0.87 for Sauria, 0.74 for Dog, 0.77 for Ruminant, 0.76 for Human, 0.76 for Rodent, and 0.72 for Aves. These values indicate strong classifier performance for Insects, Monocots, and Sauria, suggesting the model can distinguish positive and negative cases effectively in these taxa.

**Figure 5:**
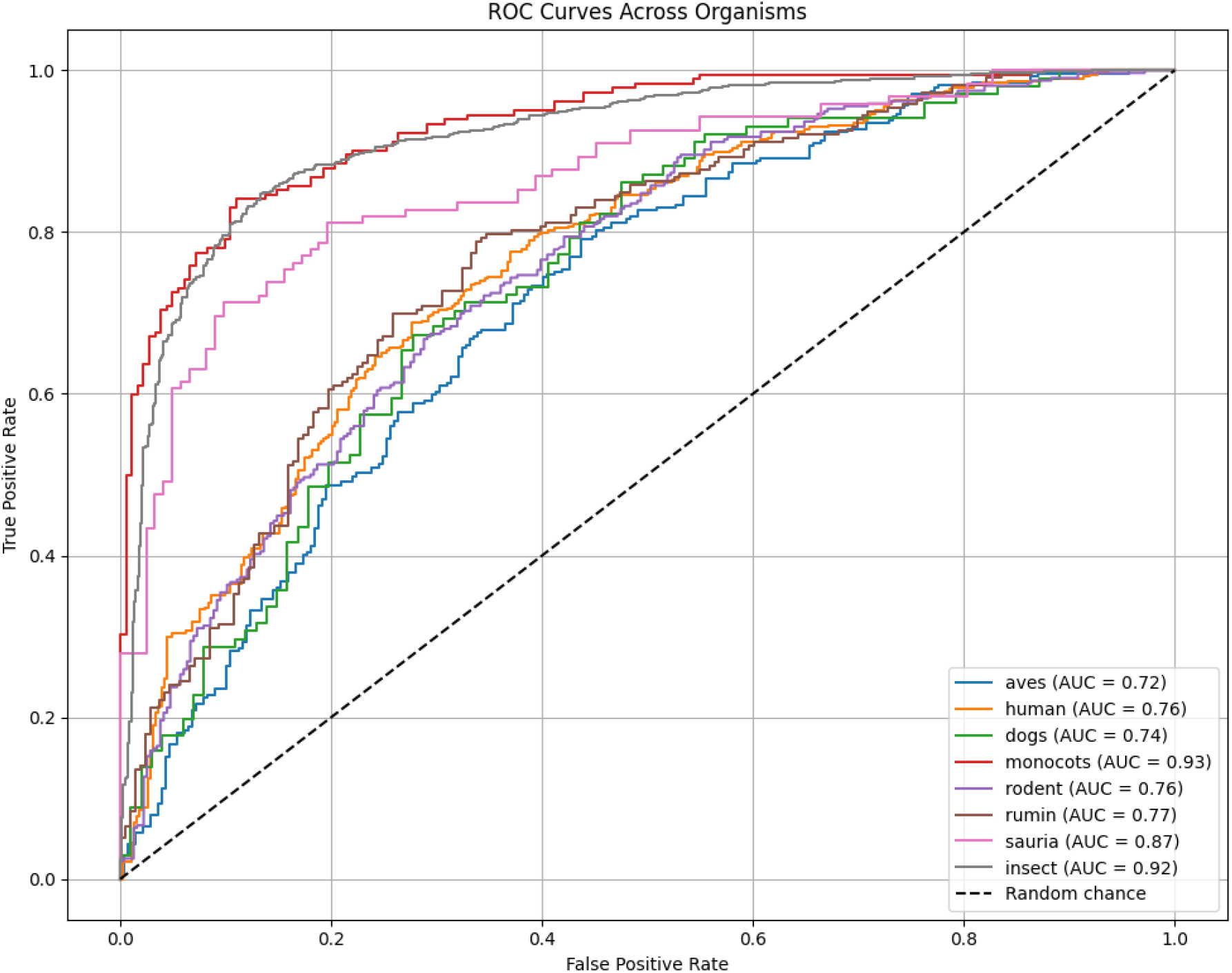
ROC-AUC of the XGBoost classifier across different organism groups. It shows how the traditional features contribute to the overall learning in different classes of organisms. Higher AUC values for Insects, Monocots, and Sauria indicate strong discriminatory performance

## 4. Discussion

### 4.1. Data collection and pre-processing

We collected all available pre-miRNA sequences of insects as our initial focus was to distinguish them from other organisms. We also collected data from rodents, monocots, aves, dogs and sauria which are reptiles. Highest number of pre-miRNA sequence from a single species was collected from humans which was followed by mouse and cattle. All the sequences formed characteristic hairpin loops which was inferred from the secondary structure calculated by RNAfold.

### 4.2. Hypothesis Testing

Our null hypothesis was that all the pre-miRNAs are physically and compositionally similar and hence performed the KS test with Bonferroni adjustment of various parameters with insects. These parameters are used in various machine learning based pre-miRNA prediction tools ^19,21,41,42^. As the p-value of parameters such as Length, GC content, MFE1, etc were < 0.05, hence we rejected the null hypothesis and accepted the alternate hypothesis that the pre-miRNA sequences from these groups vary from insects. This indicated a possibility of supervised training for classification of pre-miRNA based on their ancestral origin. Therefore, we also performed Mann Whitney and Levene’s test to check for median and variance of features among the groups. We found that all the 57 features are significant in at least on pair of organisms. Hence, we moved forward with state-of-the-art XGBoost algorithm which can efficiently learn to classify different group data based on given labels.

### 4.3. Feature engineering and model training

The estimation of features which has the maximum contribution in building the model is essential ^43,44^. Our approach initially gave us 34 features after removing highly correlated features. PCA is another widely used dimension reduction technique ^45,46^. Using PCA we selected 14 principal components as shown in the scree plot for the classification model capturing 95% of cumulative explained variance.

### 4.4. Performance evaluation

The performance of XGBoost varied among the groups. Each group had fairly good accuracy however accuracy is often misleading and cannot be considered as best indicator of performance ^47,48^. The predictive model had good specificity for each group. However, only insects, plants, and saurias had sensitivity more than 60%, with insects having the highest at 83.95%. MCC score which estimates all the parameter is a crucial indicator ^48^. The F1 score for all the organism was above 60%

But MCC of only Monocots and Insects was above 60%.

The AUC values of 0.92, 0.93 and 0.87 of insects, monocots, and reptiles (sauria) respectively, suggest that the parameters we used in XGBoost can classify these organisms.

## 5. Conclusion

In this work, we demonstrated the distinct nature of insect pre-miRNA by comparing it against that of other organisms and established that insect pre-miRNAs are significantly different from those of monocots, humans, rodents, ruminants, sauria, dogs, and aves. We further developed a predictive model using the XGBoost classifier, which effectively learned to differentiate pre-miRNA sequences across these organism classes based on a range of sequence and structural features.

In the future, this model can be implemented as a web server or standalone software tool, enabling researchers to rapidly classify unknown pre-miRNA sequences based on their likely taxonomic origin. Such a tool would be particularly valuable for annotating novel or poorly characterized genomes, assisting in evolutionary studies, and guiding experimental validation in non-model organisms. Additionally, expanding the dataset to include more taxa and incorporating deep learning-based feature extraction could further improve prediction accuracy and broaden the model’s applicability.

## 6. Data Availability

All the data used in the analysis can be found in: **https://github.com/adhiraj141092/mir_comp/tree/main/raw_data**

The files created during analysis is present in: **https://github.com/adhiraj141092/mir_comp/tree/main/processes**

## 7. Funding Agency

There was no funding for this work

## 8. Acknowledgment

The authors thank the Param-Ishan HPC facility of IIT Guwahati for providing the computational resources needed to carry out the experiments. We thank the reviewers for the valuable inputs for improving the quality of this paper.

## 9. Author Contribution

AN: Methodology, Project administration, Resources, Validation, Writing – original draft. UB: Conceptualization, Supervision, Writing – review & editing

## Reference

1. O’Brien, J., Hayder, H., Zayed, Y. & Peng, C. Overview of MicroRNA Biogenesis, Mechanisms of Actions, and Circulation. Front Endocrinol (Lausanne) 9, 402 (2018).

2. Han, J. et al. The Drosha-DGCR8 complex in primary microRNA processing. Genes Dev 18, 3016–3027 (2004).

3. Ambros, V. The functions of animal microRNAs. Nature 431, 350–355 (2004).

4. Ruvkun, G. B. The tiny RNA world. Harvey Lect 99, 1–21.

5. Cullen, B. R. Viral and cellular messenger RNA targets of viral microRNAs. Nature 457, 421– 425 (2009).

6. Ventura, A. & Jacks, T. MicroRNAs and Cancer: Short RNAs Go a Long Way. Cell 136, 586–591 (2009).

7. Zhang, Y. et al. microRNA-309 targets the Homeobox gene SIX4 and controls ovarian development in the mosquito Aedes aegypti. Proc Natl Acad Sci U S A 113, E4828–36 (2016).

8. Etebari, K. & Asgari, S. Conserved microRNA miR-8 blocks activation of the Toll pathway by upregulating Serpin 27 transcripts. 10.4161/rna.25481 10, 1356–1364 (2013).

9. Zhang, Q. et al. Genome-Wide Analysis of MicroRNAs in Relation to Pupariation in Oriental Fruit Fly. Front Physiol 10, 301 (2019).

10. Sun, X. H. et al. A novel miRNA, miR-13664, targets CpCYP314A1 to regulate deltamethrin resistance in Culex pipiens pallens. Parasitology 146, 197–205 (2019).

11. Tariq, K., Metzendorf, C., Peng, W., Sohail, S. & Zhang, H. miR-8-3p regulates mitoferrin in the testes of Bactrocera dorsalis to ensure normal spermatogenesis. Sci Rep 6, 22565 (2016).

12. Gulhane, P., Nimsarkar, P., Kharat, K. & Singh, S. Deciphering miR-520c-3p as a probable target for immunometabolism in non-small cell lung cancer using systems biology approach. Oncotarget 13, 725 (2022).

13. Song, J. et al. The microRNAs let 7 and mir 278 regulate insect metamorphosis and oogenesis by targeting the juvenile hormone early response gene Krüppel homolog. Development (Cambridge) 145, (2018).

14. Lee, C. T., Risom, T. & Strauss, W. M. Evolutionary Conservation of MicroRNA Regulatory Circuits: An Examination of MicroRNA Gene Complexity and Conserved MicroRNA-Target Interactions through Metazoan Phylogeny. https://home.liebertpub.com/dna 26, p209–218 (2007).

15. Friedman, R. C., Farh, K. K. H., Burge, C. B. & Bartel, D. P. Most mammalian mRNAs are conserved targets of microRNAs. Genome Res 19, 92–105 (2009).

16. Willmann, M. R. & Poethig, R. S. Conservation and evolution of miRNA regulatory programs in plant development. Curr Opin Plant Biol 10, 503–511 (2007).

17. Kozomara, A., Birgaoanu, M. & Griffiths-Jones, S. miRBase: from microRNA sequences to function. Nucleic Acids Res 47, D155–D162 (2019).

18. Ng, K. L. S. & Mishra, S. K. De novo SVM classification of precursor microRNAs from genomic pseudo hairpins using global and intrinsic folding measures. Bioinformatics 23, 1321–1330 (2007).

19. Batuwita, R. & Palade, V. microPred: effective classification of pre-miRNAs for human miRNA gene prediction. Bioinformatics 25, 989–995 (2009).

20. Tran, V. D. T., Tempel, S., Zerath, B., Zehraoui, F. & Tahi, F. miRBoost: boosting support vector machines for microRNA precursor classification. RNA 21, 775–785 (2015).

21. Gudyś, A., Szcześniak, M. W., Sikora, M. & Makałowska, I. HuntMi: an efficient and taxon-specific approach in pre-miRNA identification. BMC Bioinformatics 14, 83 (2013).

22. Stegmayer, G., Yones, C., Kamenetzky, L. & Milone, D. H. High Class-Imbalance in pre-miRNA Prediction: A Novel Approach Based on deepSOM. IEEE/ACM Trans Comput Biol Bioinform 14, (2017).

23. Xue, C. et al. Classification of real and pseudo microRNA precursors using local structure-sequence features and support vector machine. BMC Bioinformatics 6, 310 (2005).

24. Jiang, P. et al. MiPred: classification of real and pseudo microRNA precursors using random forest prediction model with combined features. Nucleic Acids Res 35, W339–W344 (2007).

25. Ng, K. L. S. & Mishra, S. K. De novo SVM classification of precursor microRNAs from genomic pseudo hairpins using global and intrinsic folding measures. Bioinformatics 23, 1321–1330 (2007).

26. Ye, J., Xu, M., Tian, X., Cai, S. & Zeng, S. Research advances in the detection of miRNA. J Pharm Anal 9, 217–226 (2019).

27. Condrat, C. E. et al. miRNAs as Biomarkers in Disease: Latest Findings Regarding Their Role in Diagnosis and Prognosis. Cells 2020, Vol. 9, Page 276 9, 276 (2020).

28. Huang, K.-Y., Lee, T.-Y.Teng, Y.-C. & Chang, T.-H. ViralmiR: a support-vector-machine-based method for predicting viral microRNA precursors. BMC Bioinformatics 16, S9 (2015).

29. Stegmayer, G. et al. Predicting novel microRNA: a comprehensive comparison of machine learning approaches. Brief Bioinform 20, 1607–1620 (2019).

30. Nath, A. & Bora, U. RNAinsecta: A tool for prediction of precursor microRNA in insects and search for their target in the model organism Drosophila melanogaster. PLoS One 18, e0287323 (2023).

31. Bugnon, L. A. et al. Deep Learning for the discovery of new pre-miRNAs: Helping the fight against COVID-19. Machine Learning with Applications 6, 100150 (2021).

32. Huang, K.-Y., Lee, T.-Y.Teng, Y.-C. & Chang, T.-H. ViralmiR: a support-vector-machine-based method for predicting viral microRNA precursors. BMC Bioinformatics 16, S9 (2015).

33. Jaiswal, S. et al. Development of species specific putative miRNA and its target prediction tool in wheat (Triticum aestivum L.). Sci Rep 9, 3790 (2019).

34. Fávero, L. P. & Belfiore, P. Hypotheses Tests. Data Science for Business and Decision Making 199–248 (2019) doi:10.1016/B978-0-12-811216-8.00009-4.

35. Vanhatalo, E., Kulahci, M. & Bergquist, B. On the structure of dynamic principal component analysis used in statistical process monitoring. Chemometrics and Intelligent Laboratory Systems 167, 1–11 (2017).

36. Shaharudin, S. M. & Ahmad, N. Choice of Cumulative Percentage in Principal Component Analysis for Regionalization of Peninsular Malaysia Based on the Rainfall Amount. in 216–224 (2017). doi:10.1007/978-981-10-6502-6_19.

37. Ahmad, F., Farooq, A. & Khan, M. U. G. Deep Learning Model for Pathogen Classification Using Feature Fusion and Data Augmentation. Curr Bioinform 16, 466–483 (2020).

38. Yang, H. et al. A comparison and assessment of computational method for identifying recombination hotspots in Saccharomyces cerevisiae. Brief Bioinform 21, 1568–1580 (2020).

39. Li, H. et al. dPromoter-XGBoost: Detecting promoters and strength by combining multiple descriptors and feature selection using XGBoost. Methods 204, 215–222 (2022).

40. Bi, Y. et al. An Interpretable Prediction Model for Identifying N7-Methylguanosine Sites Based on XGBoost and SHAP. Mol Ther Nucleic Acids 22, 362–372 (2020).

41. Jiang, P. et al. MiPred: classification of real and pseudo microRNA precursors using random forest prediction model with combined features. Nucleic Acids Res 35, W339–W344 (2007).

42. Kadri, S., Hinman, V. & Benos, P. V. HHMMiR: efficient de novo prediction of microRNAs using hierarchical hidden Markov models. BMC Bioinformatics 10, S35 (2009).

43. Biesiada, J. & Duch, W. Feature selection for high-dimensional data - A pearson redundancy based filter. Advances in Soft Computing 45, 242–249 (2007).

44. Saidi, R., Bouaguel, W. & Essoussi, N. Hybrid feature selection method based on the genetic algorithm and pearson correlation coefficient. Studies in Computational Intelligence 801, 3– 24 (2019).

45. Kambhatla, N. & Leen, T. K. Dimension Reduction by Local Principal Component Analysis. Neural Comput 9, 1493–1516 (1997).

46. Zhang, T. & Yang, B. Big Data Dimension Reduction Using PCA. Proceedings - 2016 IEEE International Conference on Smart Cloud, SmartCloud 2016 152–157 (2016) doi:10.1109/SMARTCLOUD.2016.33.

47. Sokolova, M., Japkowicz, N. & Szpakowicz, S. Beyond accuracy, F-score and ROC: A family of discriminant measures for performance evaluation. AAAI Workshop - Technical Report WS-06-06, 24–29 (2006).

48. Chicco, D. & Jurman, G. The advantages of the Matthews correlation coefficient (MCC) over F1 score and accuracy in binary classification evaluation. BMC Genomics 21, (2020).

